# Protein Structural Alignments From Sequence

**DOI:** 10.1101/2020.11.03.365932

**Authors:** James T. Morton, Charlie E. M. Strauss, Robert Blackwell, Daniel Berenberg, Vladimir Gligorijevic, Richard Bonneau

## Abstract

Computing sequence similarity is a fundamental task in biology, with alignment forming the basis for the annotation of genes and genomes and providing the core data structures for evolutionary analysis. Standard approaches are a mainstay of modern molecular biology and rely on variations of edit distance to obtain explicit alignments between pairs of biological sequences. However, sequence alignment algorithms struggle with remote homology tasks and cannot identify similarities between many pairs of proteins with similar structures and likely homology. Recent work suggests that using machine learning language models can improve remote homology detection. To this end, we introduce DeepBLAST, that obtains explicit alignments from residue embeddings learned from a protein language model integrated into an end-to-end differentiable alignment framework. This approach can be accelerated on the GPU architectures and outperforms conventional sequence alignment techniques in terms of both speed and accuracy when identifying structurally similar proteins.

## 1 Introduction

Proteins with unknown function can be annotated based on their similarity to proteins of known function. Protein similarity can be measured based on their sequence or their structure. A core tenet of molecular biology is that a protein’s sequence determines structure which in turn determines the protein’s biological function. If the structures are known, structural alignment is the preferred approach to align the protein residues and measure a protein similarity [1, 2, 3, 4, 5]. Unfortunately, most sequences lack a known structure. Up to now there have been on the order of 10^8^ protein sequences discovered, but only 10^5^ proteins have annotated functions [6] and only about 10^4^ of those proteins have experimentally verified structures [7].

While structural similarity would also seem to be the most direct connection to function, sequence similarity has proven to be more pragmatic. In the case where the evolutionary distance is short, sequence similarity provides a more reliable indicator of function than the hard task of interpreting molecular structure. Thus sequence alignments are the dominant sourc of protein function inference.

Conceptually, similarity is assessed by evolutionary distance through estimating the number of mutations required to transform one protein sequence to another. Algorithms such as BLAST [8], HMMER [9] and Needleman-Wunsch [10] are the state-of-the-art methods for computing sequence alignments. However, proteins with similar structures do not necessarily have similar sequences [2], resulting in the under-performance of traditional sequence alignments in remote homology tasks [11, 12].

Unfortunately, over half of all proteins do not have detectable homologs in standard sequence databases due to their distant evolutionary relationships [13]. Detecting these remote homologs would help us better understand mutagenesis [14], aid protein design [15], predict protein function [16], predict protein structure [17, 18, 19] and model evolution [20]. Moreover, these missing homologs could be critical in annotating genomes of organisms that have not been studied before. Therefore, for highly divergent genomes, structural comparison would potentially identify many functionally related proteins; which is unattainable due to the lack of structures. Thus rather than directly compare sequence to sequence, an attractive strategy is to infer structural features from each sequence and then compare those structural features. However, computing protein-structural alignments given only the protein sequence is still an outstanding challenge [21].

Recent work shows that large-scale direct structural alignment from sequence may be computationally tractable. For sufficiently small proteins *ab initio* modeling can predict protein structure from sequence. Consequently, structural alignments of laboriously generated *ab initio* predictions for genes with no recognizable sequence similarity have been aligned for multiple genomes; ergo, pure sequence-to-structure recognition. However these were computationally expensive calculations: on the order of ten-thousand CPU hours per gene [22, 23].

Desirably then, a method that directly inferred structural properties and structurally informed alignments would not only enable remote homology detection for function prediction but also accelerate those laborious structure predictions as well. [18, 19] This paper advances an approach to the grand challenge[21] of directly producing the structural alignment of two proteins from just their sequence.

Naturally, given enough training data, one can imagine many possible supervised learning approaches to this task. But as is so often the case, structurally labeled alignments are scarce due to the limited number of experimentally validated protein structures. Therefore the central objective of this paper is to validate our hypothesis that a particular unsupervised pre-training step will successfully extract structural features into a latent space. This is desirable, since there are abundantly available protein sequences for unsupervised estimation of this latent space. Our hypothesis requires that this latent space embeds more structural information than just the sequence itself carries. If true, then one can then subsequently use this embedding to enhance a supervised process over the limited set of structurally labeled data.

We validate this hypothesis by showing that the supervised training produces better structural alignments when using our unsupervised embedding in comparison to a naive sequence-only embedding. Our purpose here is not a contest against other possible methods but to determine that a language-model-derived, unsupervised, embedding carries structural features. Once that is established, then applications more sophisticated than our hypothesis validation harness become possible, and data-poor applications are made easier due to the abundance of data for training these language models.

To this end, we develop an end-to-end differentiable neural network that takes sequences as input and outputs a predicted structural alignment. This exploits transfer learning from a pre-trained protein language model that can represent a protein sequence as a set of residue embeddings. These residue-level embeddings are sequence position specific and can capture relevant structural information based on the contextual residues. The final step uses dynamic programming to estimate the expected alignment between two protein sequences, which is then fine-tuned against known structural alignments. Only recently has an efficient backpropagation through dynamic programming become practical [24]. This allows us to put together this end-to-end model for direct prediction of alignments from sequence inputs and train it on limited labeled datasets.

We show this type of model can be trained and validated on structural alignments. We further evaluate its precision and recall on a manually curated gold standard set of structures containing remote homologies that standard sequence alignment techniques such as BLAST and HMMER could not detect. As expected, the predicted alignments cannot perform as well as directly aligning known structures, but they outperform state-of-the-art sequence alignments by a sizeable margin.

### 1.1 Related Work

The most common approach to estimating sequence similarity is through computing a weighted edit distance, with weights derived to approximate the likelihood of a given mutation. Popular sequence alignment approaches often involve variants of Needleman-Wunsch [10] or Hidden Markov Models (HMM) that enable a probabilistic treatment of sequence alignment and alignment detection [25]. To limit the scope of this paper we do not discuss the many relevant works on multiple alignment [26, 27]. On the other hand, tools such as TM-align [2], Dali [3], Fast [28], and Mammoth [4], have been designed to perform protein structural alignment when protein structures are available.

Recent studies have shown the benefits of using high-capacity self-supervised protein language models to predict protein structure [11, 29, 30, 31, 32, 33, 34]. Pre-training these protein models on large unlabeled datasets can reduce the number of labeled data required to train classifiers, while improving the generalizability.

## 2 Methods

Here we build on the notion that sequence alignment algorithms should be designed to capture structural similarity for protein sequences with corresponding well defined ensemble average protein structures As shown in previous work [2], reliable alignments for many distant sequence pairs is not possible with the conventional sequence alignment algorithms. Ideally, we would like align structures instead of sequences to determine which residues are structurally analogous, and we’d like to detect cases where the sequence similarity is low and the structure similarity is high. However, many structure alignment methods aim to maximize atomic overlap between structures, ignoring potentially relevant sequence information. Here, we aim to fuse the best of both worlds, leveraging information from the protein sequence to infer structurally relevant alignments. To be able to train our model to perform optimally at the alignment end goal defined in this manner, we create a workflow that is end-to-end differentiable, with loss that is informed from the structural alignment. A major technical hurdle here is the need to perform dynamic programming with an unknown position specific alignment scoring matrix. This paper exploits a recent innovation [24] that we use to build a method that enables differentiating through the dynamic programming step, allowing training of all salient parameters and alignment scoring matrices.

Since we have only sequence, we need to capture the structural propensities of a given sequence. Many potential choices for embeddings apropos to structurally informed tasks are computationally expensive [11, 29, 30, 31, 32, 33, 34]. For this paper we have selected a well vetted language model [11] to cast the protein residues to a latent space with the intention of recovering the underlying grammars behind protein sequences.

In order to develop a model that can perform structural alignments, we propose using protein structural alignments estimated using a widely adopted structure alignment tool TM-align [2] for training. Since most protein structural alignments can be represented as linear sequence alignments [4], we argue that differentiable dynamic programming coupled with language models can enable the estimation of more complex structural alignments. The high-level workflow behind our proposed DeepBLAST algorithm is given in Figure 1

**Figure 1:**
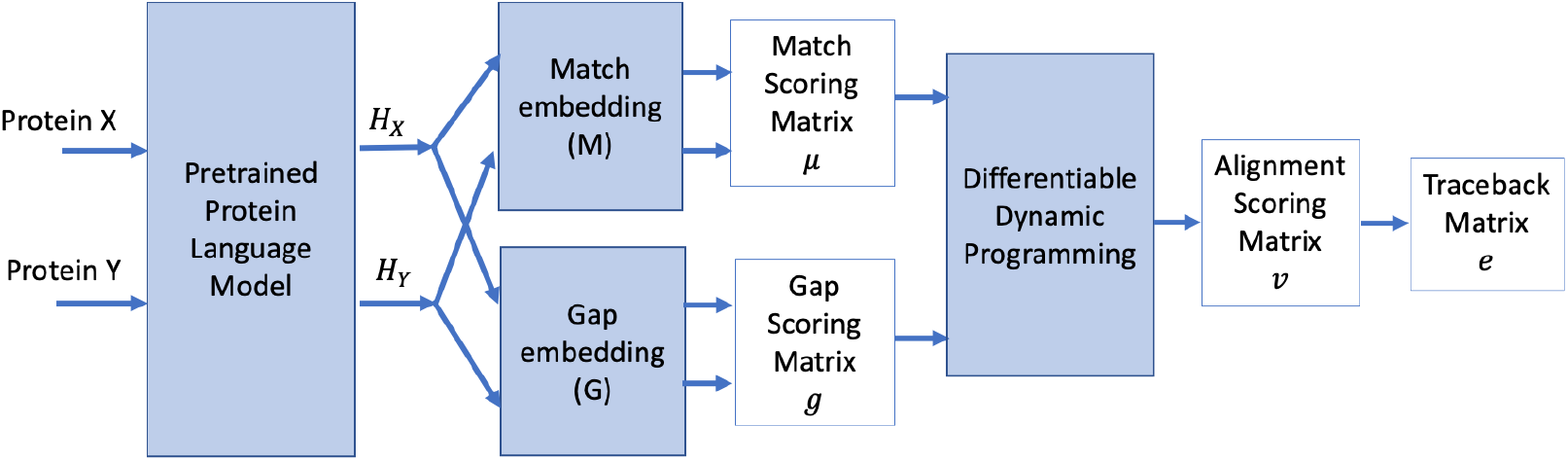
DeepBLAST workflow.

**Figure 2:**
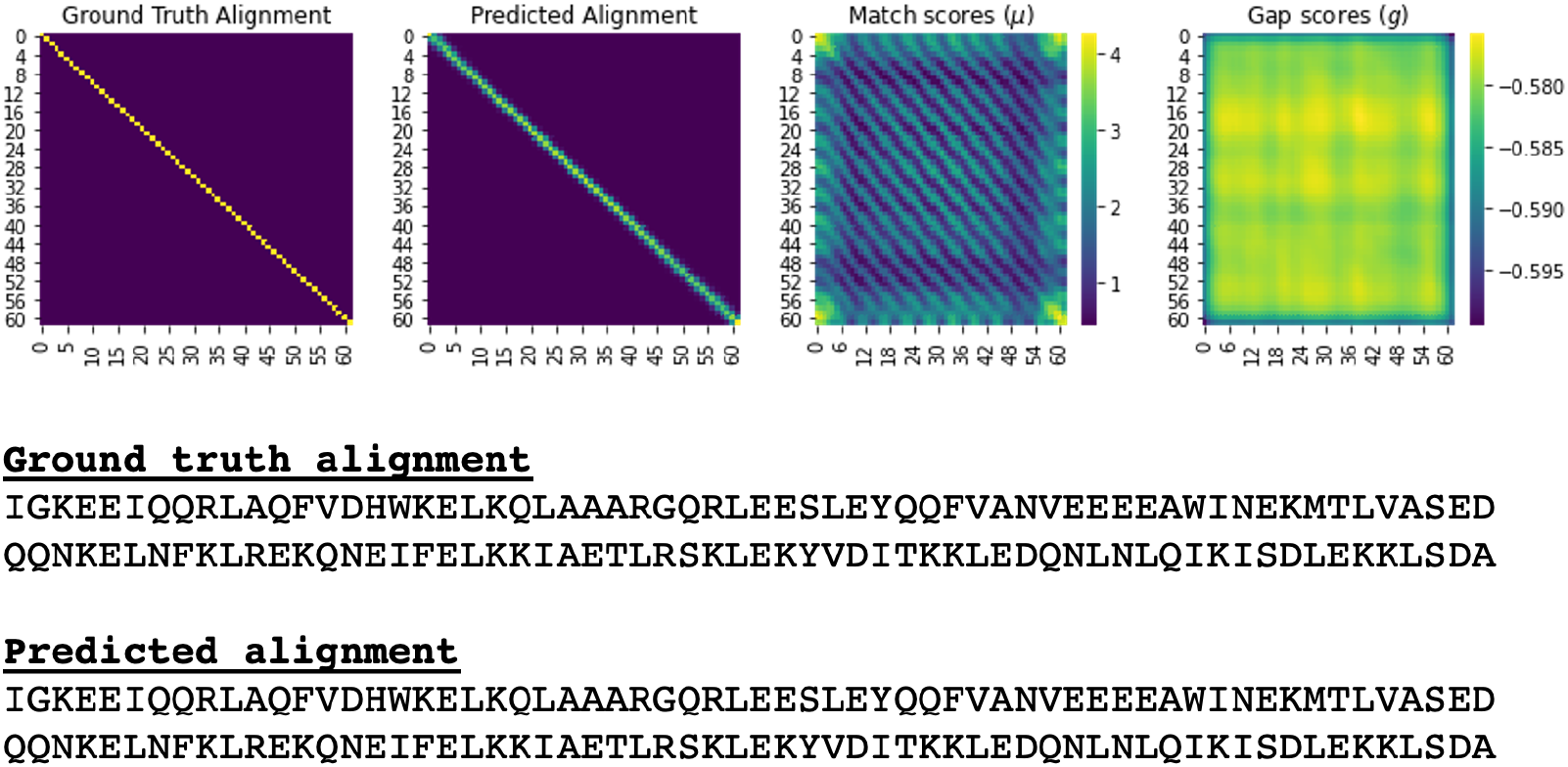
Alignment between two validation proteins: An example of an exact alignment between two proteins with very significant structural/fold similarity used to derive the ground truth alignment that has little sequence similarity.

**Figure 3:**
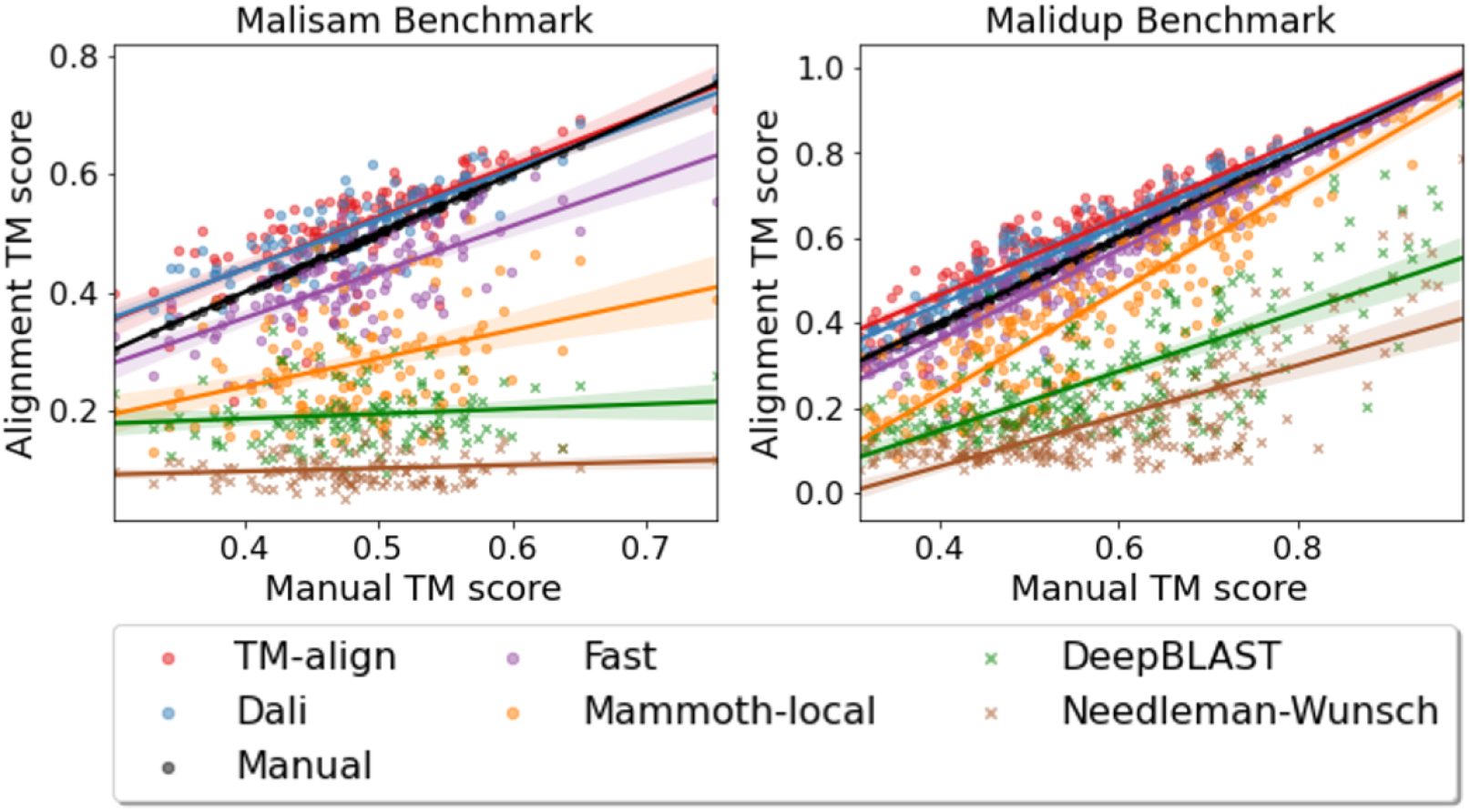
Visualization of Malisam and Malidup structural alignments. Each point represents an alignment, crosses are sequence based alignment and circles are structure based alignments. The estimated alignment is then superimposed on the structures to estimate the TM-score (Y-axis). The more to the right on the plot the more significant the structural overlap. As a reference, these are plotted against the Manual Alignment TM-score on the X-axis. The orange line derives alignment from measured secondary structure alone, while other structural alignment methods optimize a 3D structural superposition.

**Figure 4:**
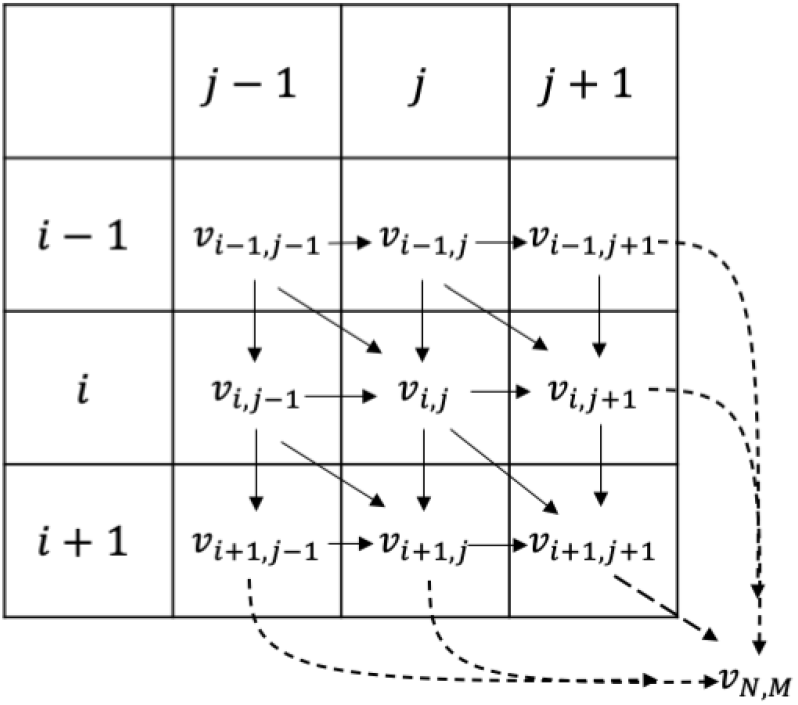
Diagram of Needleman-Wunsch recursion

Proteins *X* and *Y* are fed into the pretrained LSTM protein language model [11] to obtain embeddings *H_X_* and *H_Y_*. These residue-level embeddings are then propagated through the match embeddings (M) and gap embeddings (G) in order to obtain the match scores ***μ*** and the gap scores ***g*** as discussed in Section 2.1. The match and gap scores are used to evaluate the differentiable dynamic programming algorithm and generate a predicted alignment traceback as discussed in Section 2.2. These alignments can then be fine-tuned using a training dataset of ground truth alignments as discussed in Section 2.3 and Section 2.4.

### 2.1 Protein Language Modeling

In order to obtain an alignment from dynamic programming, the scoring parameters for matches and gaps must be obtained. We propose to utilize the pretrained protein language models to estimate these scoring parameters. These pretrained models ultimately construct a function, mapping a sequence of residues, represented as one-hot encodings, to a set of residue vectors, providing an alternative representation of these proteins. Often these models will learn these representations by being trained to predict randomly masked residues within a protein sequence. Multiple studies have shown the merits of these models when performing protein structure prediction, remote homology and protein design [30, 35, 29, 32, 31, 33, 34]. Here, we have used the pretrained LSTM PFam model from [11]. Using this pretrained language model, two proteins *X* and *Y* can be represented by embeddings ***H_X_*** ∈ ℝ^*p*×*d*^ and ***H_Y_*** ∈ ℝ^*q*×*d*^, where *p* and *q* represent the lengths of proteins *X* and *Y* and *d* is the embedding dimension of the language model. Given these representations, we can construct mappings *M* and *G* to obtain match scores and gap scores for the differentiable dynamic programming as follows

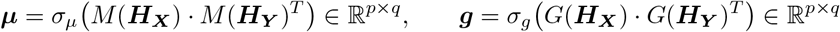

The functions *M*: ℝ^*t*×*d*^ → ℝ^*t*×*d*^ and *G*: ℝ^*t*×*d*^ → ℝ^*t*×*d*^ are intermediate functions that take in as input a set of *t* residue vectors. These functions are parameterized by LSTM networks, which can be fine-tuned through the backpropagation enabled by the differentiable dynamic programming. Activation functions *σ_μ_* and *σ_g_* are softplus and logsigmoid functions to ensure that the match scores ***μ*** are strictly positive and the gap scores ***g*** are strictly negative. These constraints are used to penalize gaps and reward matches. This also helps enforce identifiability of the model, which we have found to improve the accuracy of the model in practice.

### 2.2 Differentiable Dynamic Programming

Our proposed differential dynamic programming framework doesn’t learn any parameters; it is designed purely to enable backpropagation to fine-tune the scoring functions *M* and *G*. Differentiable dynamic programming has been extensively explored in the context of dynamic time warping [36, 24]. Koide et al [37] and Ofitserov et al [38] suggested that a differentiable Needleman-Wunsch alignment algorithm could be derived, but its implementation has remained elusive. Here, we provide the first GPU-accelerated implementation of the differentiable Needleman-Wunsch algorithm.

Previous work [24] has shown that backpropagation can be performed on dynamic programming algorithms by introducing smoothed maximum and argmax functions. Doing so will enable the computation of derivatives while providing a tight approximation to the optimal dynamic programming solution. The traditional Needleman-Wunsch algorithm can be defined with the following recursion

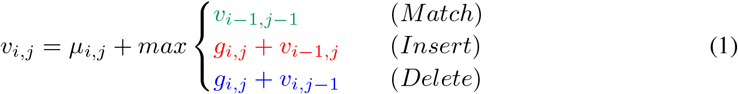

where the alignment score *v_i,j_* is evaluated on position *i* in the first sequence *X* and on position *j* in the second sequence *Y*. Sequences *X* and *Y* are of lengths *n* and *m* respectively. *μ_i,j_* represents the log-odds score of residues *X_i_* and *Y_j_* being aligned and *g_ij_* represents the log-odds score of an insertion or a deletion at positions *i* and *j*. Due to the structure of dynamic programming problems, *v_n,m_* is guaranteed to be the optimal alignment score between the two sequences. Furthermore, the optimal alignment can be obtained by tracing the highest-scoring path through the alignment matrix via argmax operations.

As neither the max nor the argmax operations are differentiable, the alignment scores and the traceback cannot be differentiated in the traditional formulation of the traceback operations needed to generate alignments. Accordingly, Mensch et al [24] introduced smoothed differentiable operators

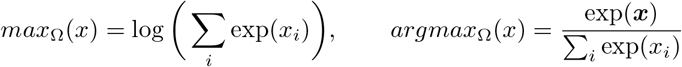

where the smooth max operator *max*_Ω_(*x*) is given by the log sum exp function and the smoothed *argmax*_Ω_(*x*) is given by the softmax function. Since the softmax function can be derived from the derivative of *max*_Ω_, the traceback matrix can also obtained by differentiating the resulting alignment matrix. The resulting traceback matrix will yield the expected alignment between the two proteins. Since the loss function is defined as the difference between the predicted traceback matrix and the ground truth traceback matrix, the derivatives of the traceback matrix also need to be defined, which requires both the computations of the directional derivatives and the local Hessians of the alignment matrix (Appendix A).

In practice, dynamic programming can be the major computational bottleneck due to GPU data transfer and the quadratic runtime of the Needleman-Wunsch algorithm. To address this, we have implemented a GPU-accelerated differentiable Needleman-Wunsch algorithm inspired by Manavski et al [39]. As can be seen from the benchmarks shown in Figure 5, this algorithm is an order of magnitude faster than the naive CPU-bound Needleman-Wunsch implementation. Furthermore, this algorithm can enable batching, allowing for multiple alignments to be processed in parallel. As shown in Figure 5, larger batch sizes can further improve the scaling over CPU-bound alignments.

**Figure 5:**
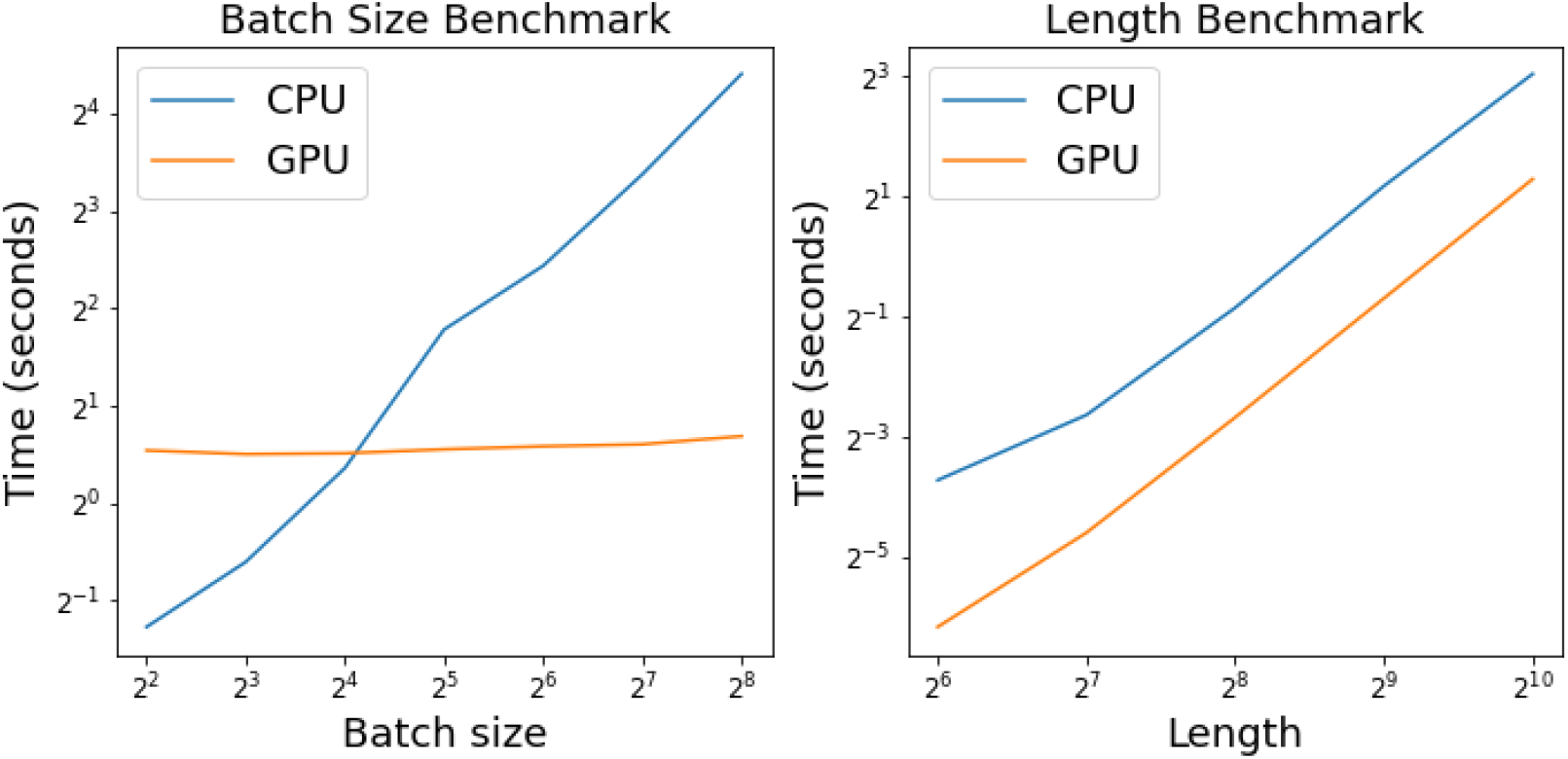
CPU vs GPU Differentiable Needleman-Wunsch benchmarks. The batch-size benchmark was run with randomized proteins of length 800 and the length benchmark was run with a fixed batch size of 64.

### 2.3 Alignment Loss Function

By defining a loss function between the predicted alignment and the structural alignment from TM-align, we can evaluate the accuracy of DeepBLAST and fine-tune the functions *M* and *G*. Mensch et al [24] proposed using the Euclidean distance between the predicted and ground truth alignments as a loss function. In practice, we found that a cross-entropy loss provided more reasonable alignment results. This loss is given by

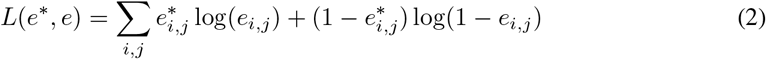

where *e*^*^ is the ground truth alignment and *e* is the predicted alignment. As shown in [24], the predicted traceback matrix represents the expectation across all possible predicted alignments, which is represented as a matrix of probabilities. As a result, the resulting alignment problem can be interpreted as a classification task of identifying whether two residues between a pair of proteins are alignable. This provides additional motivation for utilizing cross-entropy as a loss function.

### 2.4 Training

We trained DeepBLAST on 1.5M alignments from the PDB [40] obtained using TM-align [41]. These proteins were obtained from a curated collection of 40k protein structures [42]. Details behind the model specification and training can be found in Appendix C.

## 3 Results

### 3.1 Assessing alignment quality via held out analysis

Alignment accuracy was assessed on a held out test dataset of 79k structural alignments. To determine how well DeepBLAST generalizes, proteins that were in both the heldout LSTM PFam dataset [11] and the held out TM-align alignments used to train DeepBLAST were analyzed. Within the DeepBLAST held out dataset, 57444 alignments were constructed from proteins that were unique to the DeepBLAST held out dataset, 1853 alignments contained proteins that were similar to those trained from the LSTM PFam training dataset and 19967 alignments contained a single protein that was unique to the DeepBLAST held out dataset and a single protein that was in the LSTM PFam training dataset. To evaluate the accuracy of the alignments, precision and recall were computed from the number of correctly identified matching residues. Since each alignment can be represented as a bipartite graph where the edges represents matching residues between two proteins, precision and recall can be extracted from comparing the edge sets between the predicted alignment and the ground truth alignments. Figure 6 shows the distribution of correctly identified alignment edges.

Within the DeepBLAST held out dataset, the true positive distribution of proteins held out from the training roughly resembles the true positive distribution of proteins observed in pre-training. The average true positive rate, false positive rate and false discovery rate are shown in Table 2.

**Figure 6:**
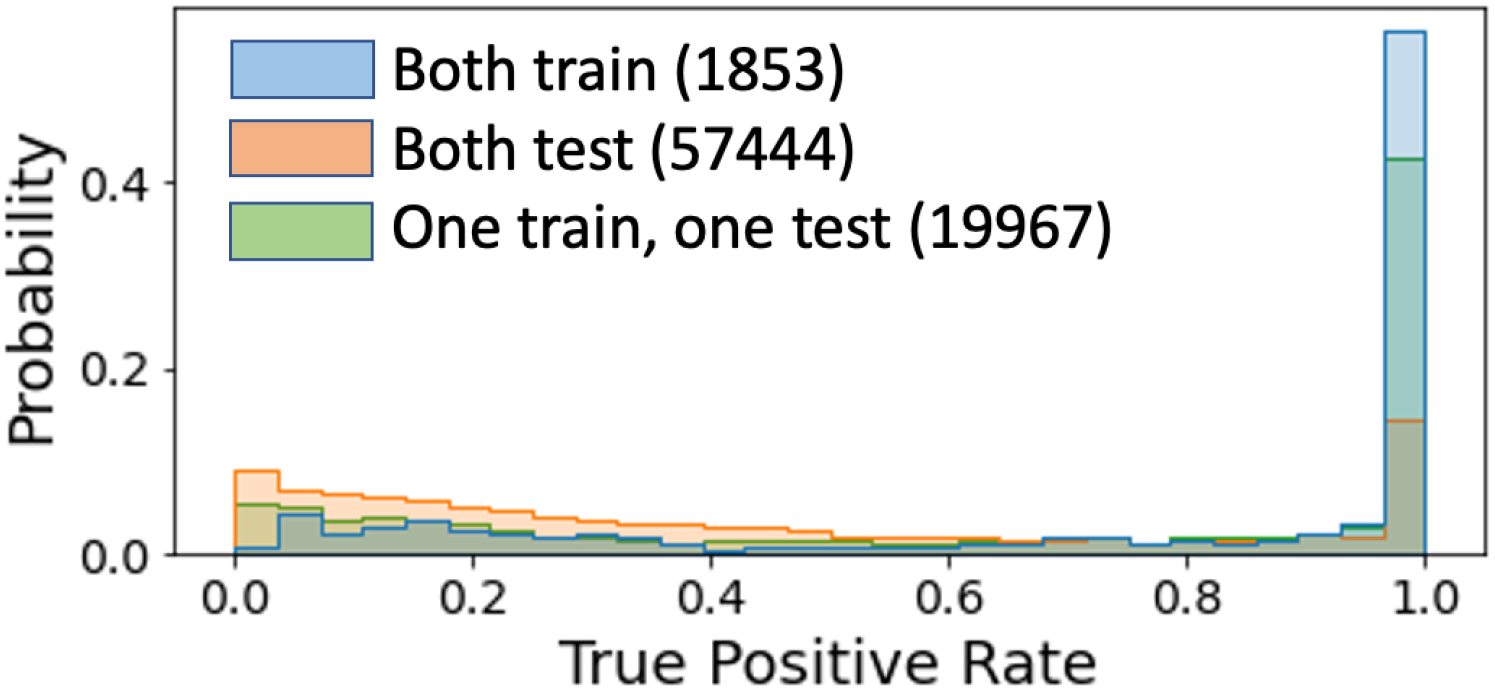
Distribution of DeepBLAST true positive rates across heldout alignment datasets.

**Figure 7:**
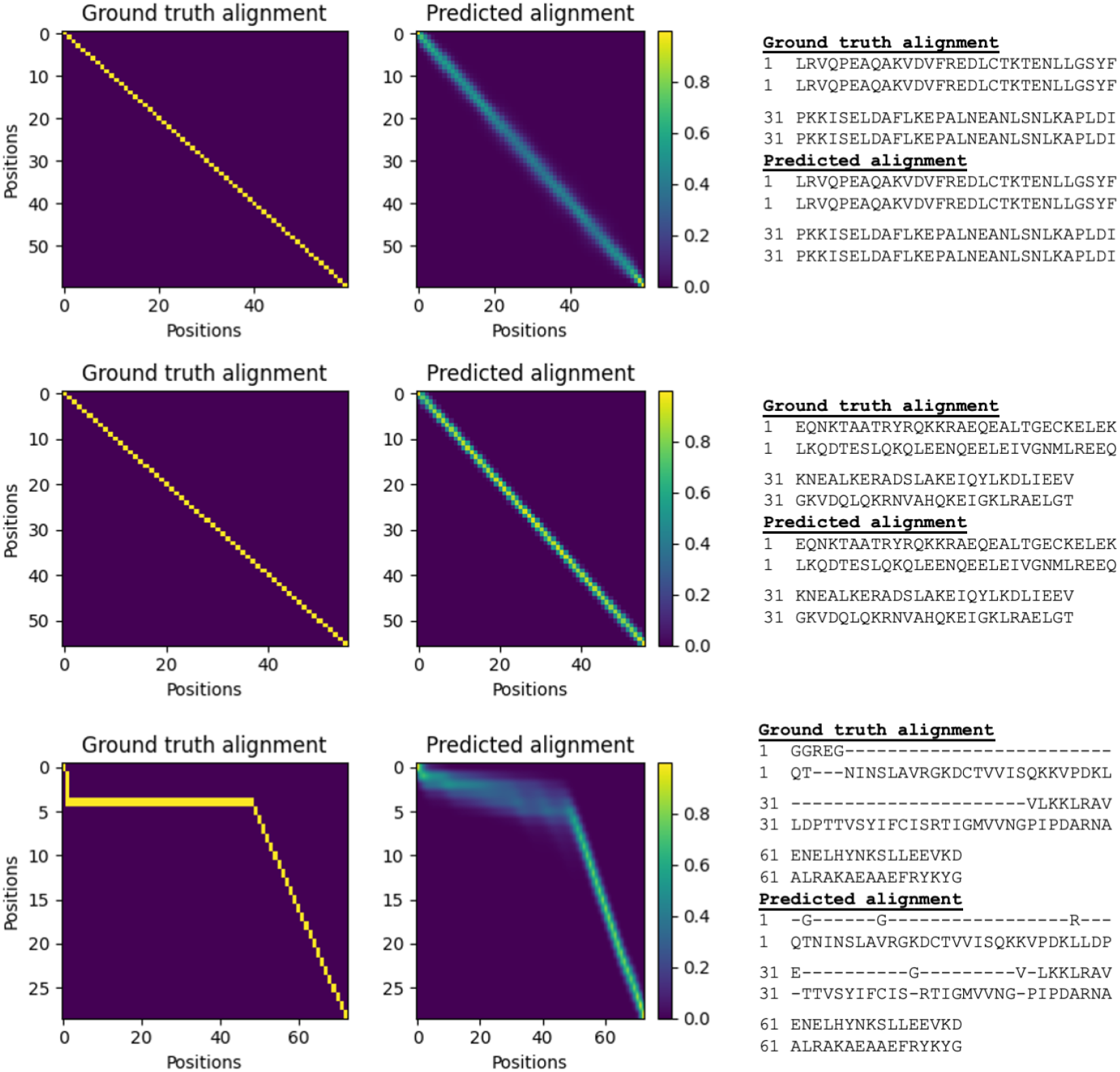
Examples of predicted alignments on the validation dataset.

As expected, DeepBLAST performs best with the TM-align structural alignments on sequences that have been used for training the LSTM language model. This is observed in the true positive rate in addition to the false positive and false negative rates, as shown in Table 2. Thus, it appears that the generalization of DeepBLAST primarily hinges on the underlying language model, as suggested by Rao et al [31].

### 3.2 Manually Curated Structural Alignment Benchmarks

We benchmarked DeepBLAST against three sequence alignment methods, Needleman-Wunsch, BLAST and HMMER in addition to four structural alignment methods that work directly with the atomic coordinates, namely FAST, TM-align, Dali and Mammoth-local. TM-align emphasizes achieving the simultaneous maximal 3D spatial overlap of the atoms in each protein. Conversely, the local structure alignment scores feasible residue pairings between the proteins according to structural similarity of just seven-contiguous-neighbor windows, apropos to a remote homology philosophy where the full length structure is allowed to be flexible and so does not require all the aligned atoms to overlap simultaneously after a rigid body orientation. Dali utilizes distance matrix computed from hexapeptide contacts to align the two protein structures. FAST tries to preserve similar residue-residue contact patterns. We extracted the local structure alignment from first phase of the Mammoth algorithm.

Thus from emphasizing long-range overlap, contacts, and local-window similarity, these reference algorithms span the rational disagreement across different expert opinions for the most meaningful structure alignment considering only backbone atomic coordinates (C-alpha or C-beta atoms). All of those algorithms disregard sequence similarity. Our method uses sequence alone; we do not supply the atomic coordinates of either protein to the algorithm after training it.

To form a common reference for all nine definitions of the optimal alignment, we designated a gold standard to be the manually curated structural alignments. Manual structure alignment is intuitive human assessment typically emphasizing 3D overlap and topology preservation since those features are easier to visualize than a plethora of local alignments and contacts [43, 44, 45].

All methods tend to agree when the problem is trivial due to near sequence identity and thus near structural identity. Therefore the most valuable gold-standard is where the dataset members have low sequence identity as well as varied degrees of structural similarity. In that regime, human intuition can provide an informative baseline by accessing additional evolutionary knowledge. Our benchmarks were performed on the curated Malisam [46] and Malidup [47] protein structural alignment benchmarking datasets.

As shown in Table 1, we observe that DeepBLAST outperforms all of the sequence alignment models by a large margin. This is observed in terms of both precision and recall as shown in Figure 8. In both benchmarks, the sequence similarity between proteins was below the observed detection limit for both BLAST and HMMER. As a result, these tools were not able to detect the vast majority of the alignments. This leaves Needleman-Wunsch as the baseline for sequence alignment methods.

**Table 1:** Malisam and Malidup Benchmarks. Sequence and structure alignment methods measured by their F1 score. Fast, TM-align, Dali Mammoth-local are structure-structure alignment methods and provide an structure-informed upper bound for this benchmark, as many of the most challenging alignments in this benchmark are ultimately structure-derived or curated with a structure-structure alignment as an oracle.

|  | Malidup |  | Malisam |  |
| --- | --- | --- | --- | --- |
| Method | F1 Score | # Detected | F1 Score | # Detected |
| BLAST | $0.019 \pm 0.019$ | 5 | $0.000 \pm 0.000$ | 2 |
| HMMER | $0.020 \pm 0.02$ | 8 | $0.020 \pm 0.020$ | 3 |
| Needleman-Wunsch | $0.098 \pm 0.010$ | 234 | $0.025 \pm 0.003$ | 129 |
| DeepBLAST | $0.144 \pm 0.012$ | 234 | $0.063 \pm 0.007$ | 129 |
| Mammoth-local | $0.483 \pm 0.020$ | 234 | $0.187 \pm 0.017$ | 129 |
| Fast | $0.569 \pm 0.026$ | 234 | $0.300 \pm 0.030$ | 129 |
| TM-align | $0.576 \pm 0.024$ | 234 | $0.393 \pm 0.031$ | 129 |
| Dali | $0.791 \pm 0.014$ | 234 | $0.619 \pm 0.029$ | 129 |

**Table 2:** Breakdown of the DeepBLAST True positive, False positive and False negative rates on the Held out alignment datasets.

| Dataset | TPR | FNR | FDR |
| --- | --- | --- | --- |
| Both test | 0.421 | 0.579 | 0.584 |
| One test, one train | 0.649 | 0.350 | 0.351 |
| Both train | 0.755 | 0.245 | 0.248 |

**Figure 8:**
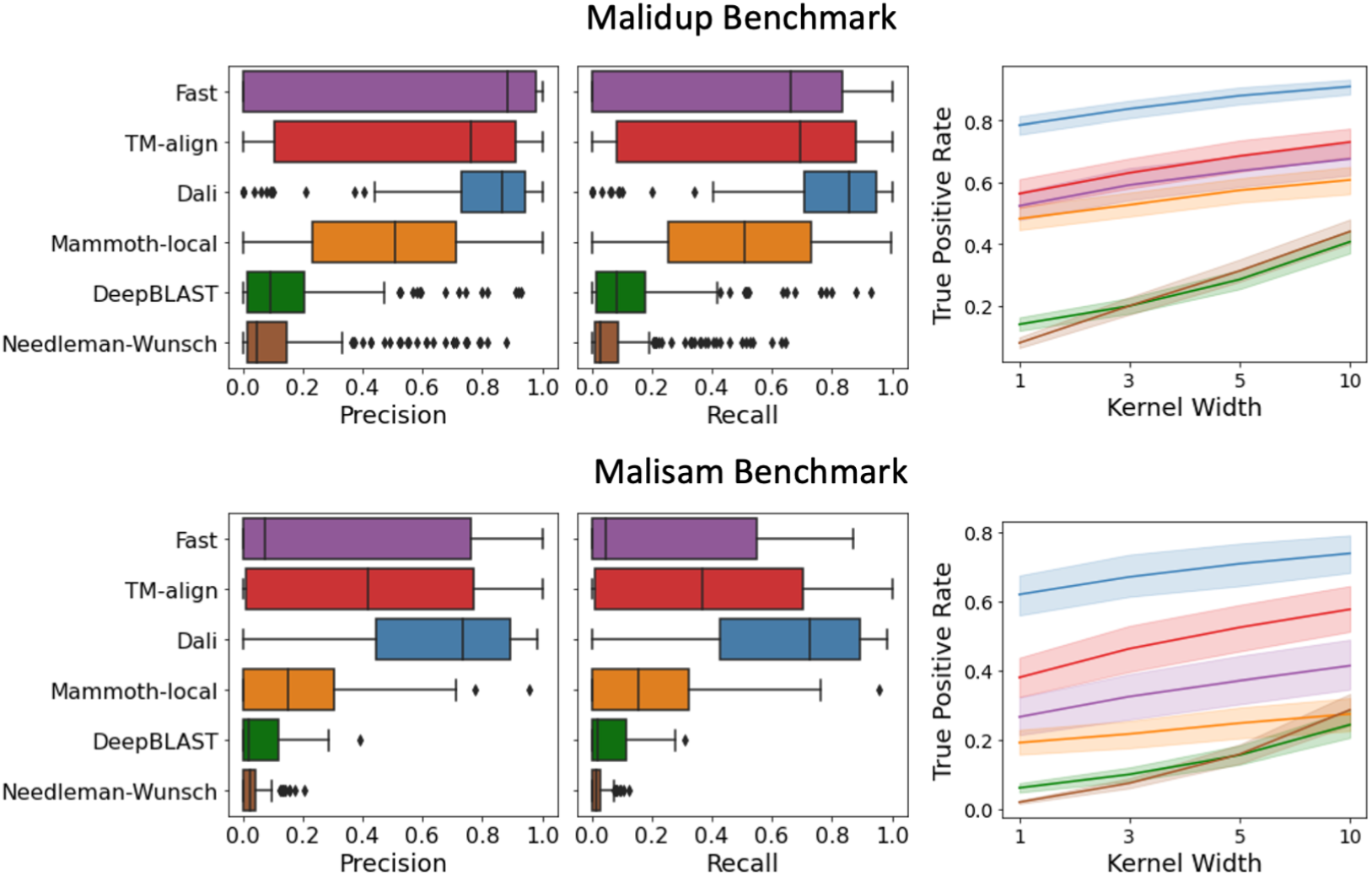
Precision and Recall metrics for each alignment on Malidup and Malisam benchmarks. The true positive rate was evaluated within window.

However, there is no one definition of what the best structural alignment is [25, 48]. This task becomes increasingly ambiguous as the remoteness of the homolog increases and the number of homologous residues declines. Thus two sequence alignments might slightly disagree but still be equally good in terms of structural superposition. Thus the above F1 score is indicative of alignment accuracy but is rigid since it only scores sequence alignments exactly matching the reference.

A better measure than the true positive rate is to directly measure the degree of structural overlap of two proteins given a specified alignment. The TM-score is one calibrated measure of this that factors out the dependence of the number of partially superimposed residues on the length of the protein. Figure 3 displays the TM-scores for multiple accepted criteria for superimpose 3D structures (Dali, TM-align, Fast). Each is plotted against the TM-score of a human curated manual superposition. The scatter in these points represents the reasonable disagreement among these varied structural similarity criteria, since all are arguably good methods. One can even observe that TM-align and Dali actually achieve slightly higher TM-scores than the supposedly ideal manual curation, highlighting the uncertainty in the best structural superposition. It can be seen that all of the structure aware methods agree at high structural similarity, TM-score=1 being perfect superposition of all atoms, but disagree increasingly as the TM-score declines.

To determine the agreement between sequence alignment methods and direct alignment by known structure, the TM-score was calculated for the predicted alignment. Both sequence alignment methods under perform the structure aware methods in terms of their TM-scores. However it is apparent that DeepBLAST is nearly always able to superimpose structures better than Needleman-Wunsch. We also compared these to the scores generated by using just local secondary structure to perform the alignment. This is essentially a Needleman-Wunsch alignment using local structural similarity to determine the quality of the alignment based on the structures.

As shown in Figure 3, DeepBLAST is competitive with Mammoth-local while consistently outperforming Needleman-Wunsch. We suspect that a large part of the disagreement between DeepBLAST and the structural alignment methods could be explained by the different strategies towards handling gaps. DeepBLAST does not have a mechanism that handles affine gaps, which would be expected to cause it to over-align sequence instead of allowing large gaps for insertions or deletions. Figure 7 showcases a DeepBLAST predicted alignment with an affine gap; the long gaps are over-penalized and DeepBLAST is forced to insert intermediate matches in order to obtain an optimal alignment. This is more apparent from the TM-scores and RMS values highlighted in Figure 9; these metrics suggest that DeepBLAST currently has difficulties obtaining accurate global structural alignments, which may be partially attributed to the inability to handle affine gaps. Handling affine alignments in a differentiable dynamic programming framework is currently an outstanding problem and we discuss potential approaches to adding afine gaps, as an area for future work, below. In spite of this shortcoming, there is evidence that DeepBLAST is able to learn structural information from the sequence. From the PSI scores shown in Figure 9, the high confidence alignments predicted by DeepBLAST are largely in agreement with the manually curated structural alignments. Furthermore, the sequence identity scores in Figure 9 reveal that DeepBLAST is able to obtain structural alignments that have less than 25% sequence identity, a known barrier for sequence alignment methods but can be resolved with the known protein structures. All together, these metrics suggest that DeepBLAST can perform local structure alignment.

**Figure 9:**
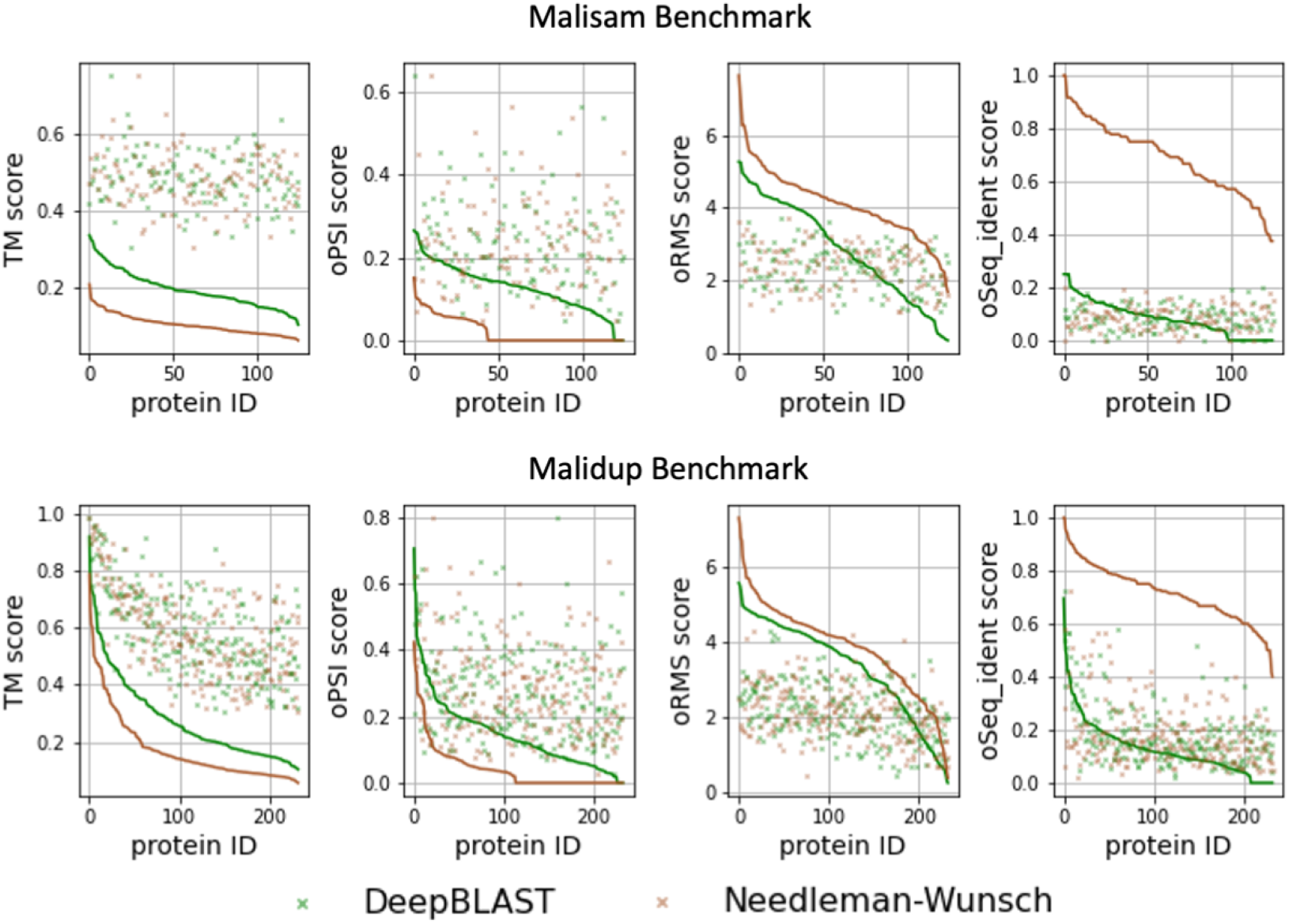
Comparison of DeepBLAST and Needleman-Wunsch on the Malidsam and Malidup benchmark. TM score measures the superposition agreement between the two aligned protein structures. The oPSI metric measures the fraction aligned residues relative to the smaller protein on the aligned residues predicted to strongly superimposed by the alignment method. The oRMS metric measures the root mean standard deviation of the atomic positions on the aligned residues predicted to strongly superimposed by the alignment method. The oSeq identity score measures the fraction of identical sequence measured over the subset of the sequence alignment that was also aligned structurally by method. All of the alignment metrics are displayed in rank order, and the points represent the manual scores for that given protein, representing and upper or lower bound of the correct alignment.

## Conclusion

The major finding of this work is that language model embeddings capture much more of the structural basis for alignment than a purely sequence based alignment when used in an end to end differentiable framework with a structure-alignment based loss. This shows that the objective function being optimized by our method is strongly correlated to the desired objective of capturing the structural basis for alignment that is implicit in the sequence. Our findings show that our proposed method can generalize well on sequences observed by the protein language model and can align sequences where there is little local sequence similarity.

As mentioned above, the spread in the true positive rate for the 4 gold standard structure based alignment should not interpreted as performance differences across structural alignment methods.

Instead, it reveals the widespread disagreement between experts regarding the ground truth structural alignment. While sequence-only DeepBLAST does not perform well compared to these explicit structural alignment methods, the resulting DeepBLAST alignments do agree more with the structural alignment methods than the other sequence-only methods.

One major difference between Needleman-Wunsch and our proposed DeepBLAST algorithm is we are not mainly weighting the alignment according to evolutionary distance. The embeddings learned from DeepBLAST are able to capture position-specific structural hints in the sequence. Given that protein secondary structure and tertiary contacts can be predicted from sequence alone [49, 50, 51], this not surprising. Furthermore, DeepBLAST can be more readily scalable to large proteins, potentially enabling structural similarity search where *ab initio* models cannot be built. This is key for enabling more accurate sequence search, since the vast majority of protein sequences do not have known protein structures.

As is stands DeepBLAST is already a better alternative to traditional sequence similarity alignment, and is usable as such. Moreover, having validated the hypothesis that the language-model unsupervised training embeds structural attributes we can refine and adapt this signal for future applications, such as function prediction and protein structure prediction. Conveniently, the end-to-end differentiable design facilitates retraining for each new objective.

The method seamlessly and continuously bridges both the sequence clues and the structural inferences. What this paper establishes is that this combined sequence/structure feature space exists, can be learned from language models, and is robust on held-out data. Our confidence in this conclusion is high because this signal was found even in gold standards chosen for resistance to sequence alignment.

## Code Availability

Our software and analyses can be found on Zenodo at 10.5281/zenodo.4117030.

## Acknowledgements

We would like to thank Ian Fisk and Nick Carriero for providing the computer support required to train these models. We also want to thank Strategic Blue for providing funding for benchmarking. We want to acknowledge Michael Heinzinger, Ahmedand Elnaggar, Christian Dallago, Konstantin Weibenow from TU Munich, Prathima Srinivas and Emily Webber from AWS, and Rob Knight and Igor Sfiligor from UCSD for their discussions. CEMS thanks LANL LDRD project 20200222DR (XX4200) for support. We want to acknowledge Pytorch [52], Pytorch-Lightning [53], Biopython [54], Matplotlib [55], Scipy [56] and Numpy [57] for providing the software foundation that this work was built upon.

## Appendices

## A Differentiable Needleman-Wunsch Algorithm

Recall from Equation 1 the recursion behind Needleman-Wunsch is given by

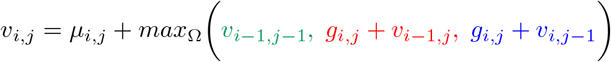

where 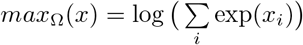 is the smoothed maximum operator. The recursion scheme can be visualized as shown in Figure 4

If one interprets the Needleman-Wunsch algorithm as a special case of a paired Hidden Markov Model [25], then the smoothed maximum operator exactly corresponds to the aggregation operator in the Forward algorithm. Given 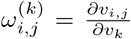 for *k* ∈ {(*i* + 1*, j*), (*i, j* + 1), (*i* + 1*, j* + 1)} the derivative of the terminal forward score *v_N,M_* can be obtained via

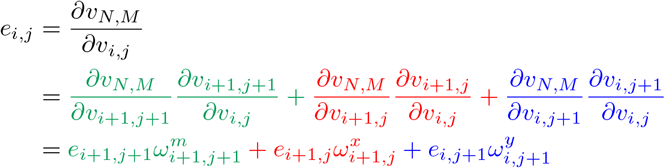

This derivative formulation will also enable backprogation to downstream parameters. Letting 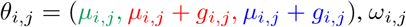 can also be given by 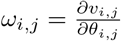.

The computation of the alignment scores and traceback matrices for the differentiable Needleman-Wunsch algorithm is given in Algorithm 1. Insertion states (*x*), match states (*m*) and deletion states (*y*) have all been color coded red, green and blue respectively. As proposed by Mensch et al, *e_i,j_* can be interpreted as elements in an expected alignment. With this in mind, the expected alignment can be compared to the ground truth alignment to estimate the loss as highlighted in Equation 2. Performing backpropagation on this loss function requires the computation of local Hessians on *v_N,M_*. We will refrain from providing the derivation of these Hessians, but the computation of the derivatives of *v_i,j_, ω_i,j_, e_i,j_* are denoted by 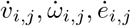, which can be computed as shown in Algorithm 2. These Hessians are calculated from directional derivatives that depend on ***Z***, the gradient of the binary cross entropy loss with respect to the variables *μ* and *g*.

It is important to note that all of the differentiable dynamic programming is only required for training. When performing prediction, the standard Needleman-Wunsch algorithm can run on the learned scoring matrices *μ* and *g*.

**Algorithm 1.**
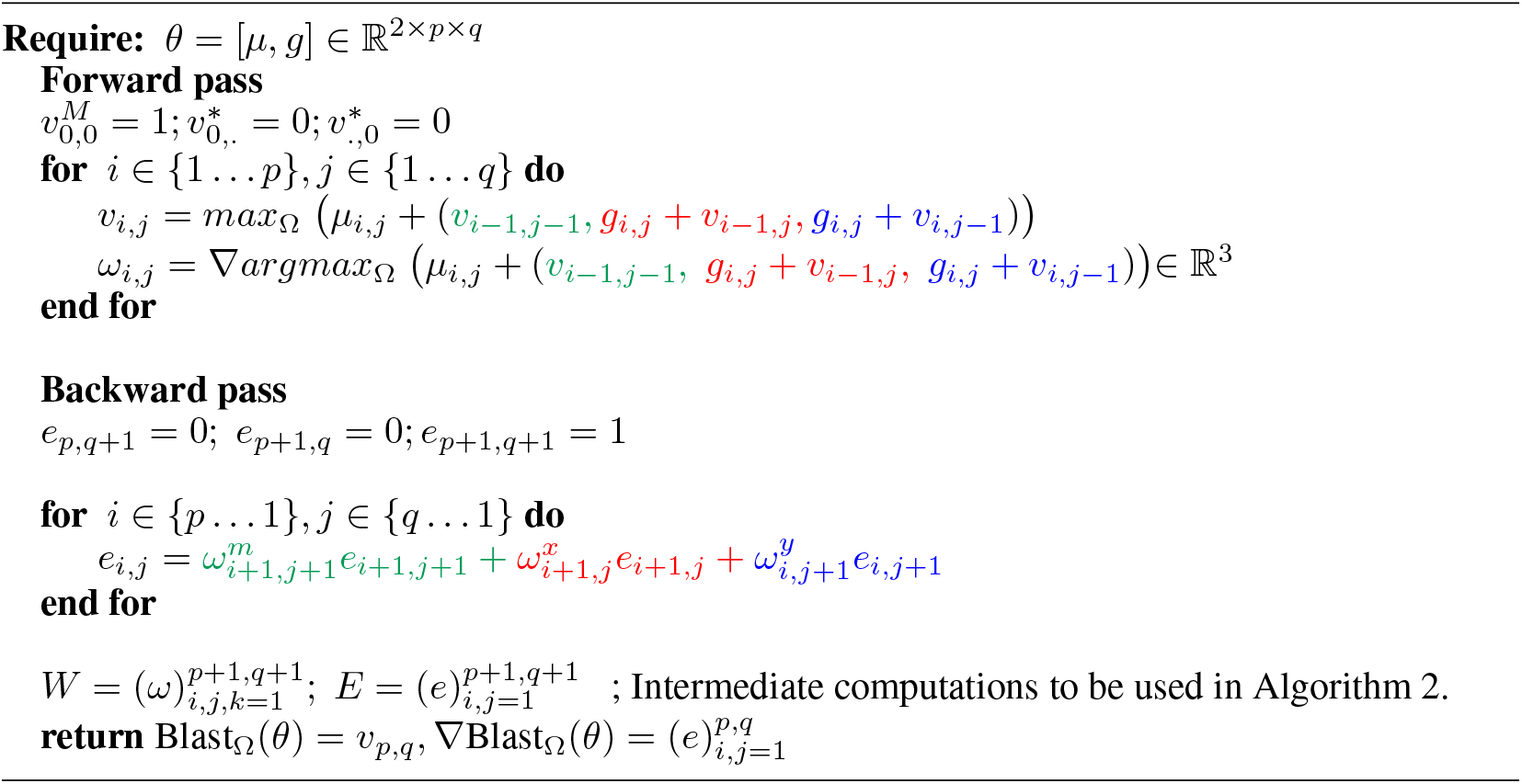
Compute Blast_Ω_(*θ*) and ∇Blast_Ω_(*θ*)

**Algorithm 2.**
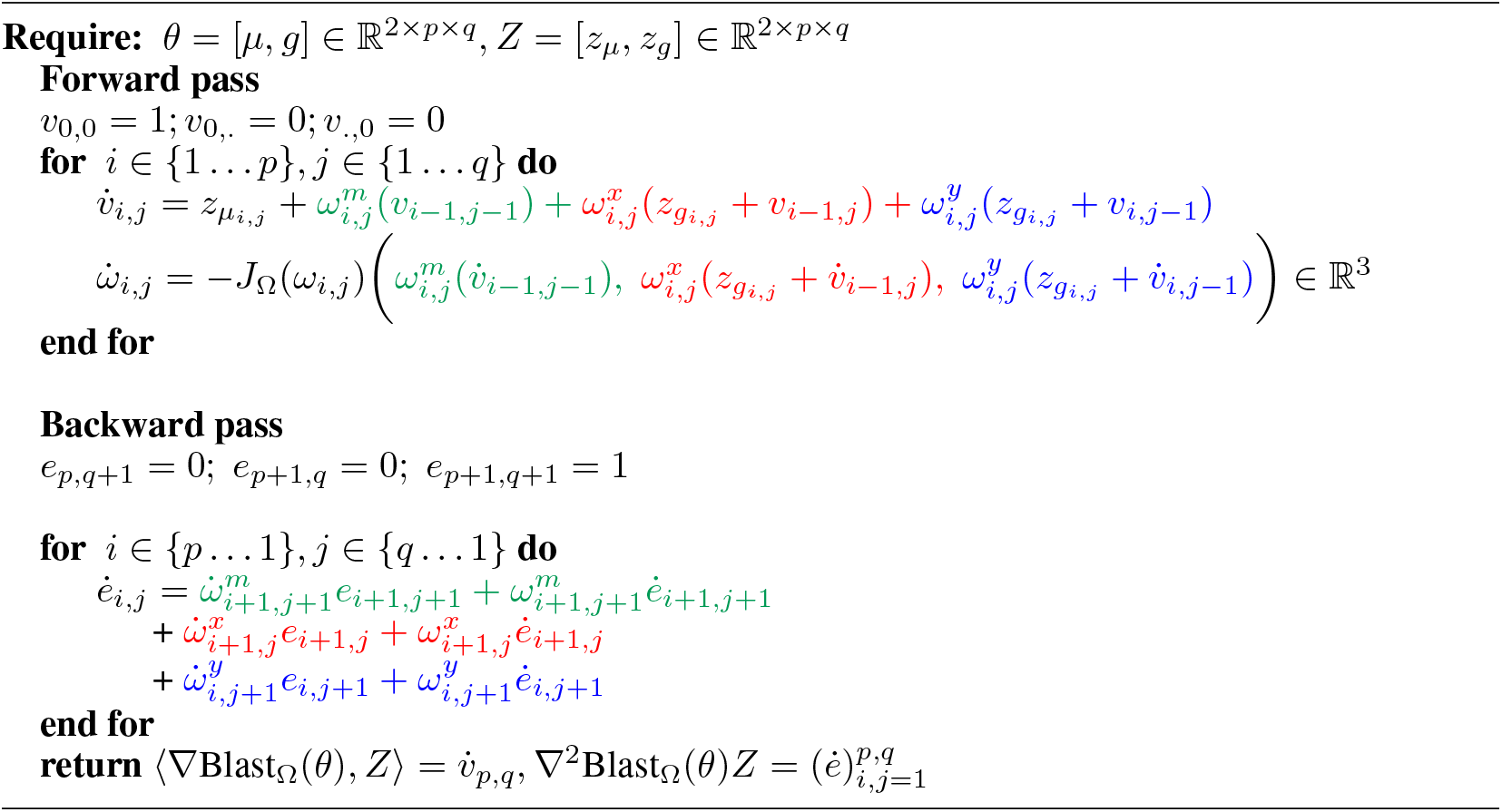
Compute (∇Blast_Ω_(*θ*)*, Z*) and ∇^2^Blast_Ω_(*θ*)*Z*

## B Runtime benchmarks

## C Additional Training Details and Held out Analysis

The final DeepBLAST model consisted of 4 LSTM layers of dimension 512 to parameterize the match embeddings *M* and gap embeddings *G*. A 2 layer bidirectional LSTM protein language model pretrained by [11] was used as a precursor step for estimating residue vectors. The resulting model had a total of 100M parameters. We used the ADAM optimizer to train the weights with an initial learning rate of 5 × 10^−5^ and the pretrained LSTM model weights were frozen. A batch size of 160 alignments was used for training. DeepBLAST was trained for 10 epochs on 4 Nvidia V100 GPUs for 4 days.

The training dataset consisted of proteins from the PDB [7]. Only proteins that had less than 1000 residues and alignments with a TM-score greater than 0.4 were considered. Furthermore, since only global alignments can be handled, the gaps at the ends of the alignment were trimmed before training. The data was split into 80/10/10 train/validation/test splits.

To evaluate how well DeepBLAST can generalize across unobserved data, the alignment accuracy was evaluated on the DeepBLAST heldout testing dataset. The intersection between the held out PFAM test sequences used to train the LSTM and the heldout TM-align alignments were determined by performing a pairwise alignment between the two held out datasets with BLAST. Alignments whose sequences were both that were detected to be homologous to sequences in the hold out PFAM dataset according to BLAST are labeled as “Both train”. Sequences that were only one sequence was found in the hold out PFam dataset are labeled as “One test, one train”. Sequences where neither sequence was found in the PFam dataset was labeled as “Both test”. As shown in Table 2, DeepBLAST generalizes best on sequences observed in the LSTM pretraining procedure.

## Notes

### Competing Interest Statement

The authors have declared no competing interest.

## References

[1] Jinbo Xu, Ming Li, Dongsup Kim, and Ying Xu. Raptor: optimal protein threading by linear programming. Journal of bioinformatics and computational biology, 1(01):95–117, 2003.

[2] Yang Zhang and Jeffrey Skolnick. Tm-align: a protein structure alignment algorithm based on the tm-score. Nucleic acids research, 33(7):2302–2309, 2005.

[3] Liisa Holm, Sakari Kääriäinen, Chris Wilton, and Dariusz Plewczynski. Using Dali for Structural Comparison of Proteins. Current Protocols in Bioinformatics, pages 1–24, 2006.

[4] Angel R. Ortiz, Charlie E.M. Strauss, and Osvaldo Olmea. MAMMOTH (Matching molecular models obtained from theory): An automated method for model comparison. Protein Science, 11(11):2606–2621, 2009.

[5] Vladimir Gligorijevic, P Douglas Renfrew, Tomasz Kosciolek, Julia Koehler Leman, Kyunghyun Cho, Tommi Vatanen, Daniel Berenberg, Bryn C Taylor, Ian M Fisk, Ramnik J Xavier, et al. Structure-based function prediction using graph convolutional networks. bioRxiv, page 786236, 2019.

[6] Amos Bairoch and Rolf Apweiler. The swiss-prot protein sequence database and its supplement trembl in 2000. Nucleic acids research, 28(1):45–48, 2000.

[7] Gary Gilliland, Helen M. Berman, Helge Weissig, Ilya N. Shindyalov, John Westbrook, Philip E. Bourne, T. N. Bhat, and Zukang Feng. The Protein Data Bank. Nucleic Acids Research, 28(1):235–242, 01 2000.

[8] Stephen F Altschul, Warren Gish, Webb Miller, Eugene W Myers, and David J Lipman. Basic local alignment search tool. Journal of molecular biology, 215(3):403–410, 1990.

[9] Robert D Finn, Jody Clements, and Sean R Eddy. Hmmer web server: interactive sequence similarity searching. Nucleic acids research, 39(suppl_2):W29–W37, 2011.

[10] Saul B Needleman and Christian D Wunsch. A general method applicable to the search for similarities in the amino acid sequence of two proteins. Journal of molecular biology, 48(3):443–453, 1970.

[11] Tristan Bepler and Bonnie Berger. Learning protein sequence embeddings using information from structure. 7th International Conference on Learning Representations, ICLR 2019, pages 1–17, 2019.

[12] Shumin Li, Junjie Chen, and Bin Liu. Protein remote homology detection based on bidirectional long short-term memory. BMC bioinformatics, 18(1):1–8, 2017.

[13] Chiara Vanni, Matthew S Schechter, Silvia Acinas, Albert Barberán, Pier Luigi Buttigieg, Emilio O Casamayor, Tom O Delmont, Carlos M Duarte, A Murat Eren, Robert D Finn, et al. Light into the darkness: Unifying the known and unknown coding sequence space in microbiome analyses. BioRxiv, 2020.

[14] Ivan Adzhubei, Daniel M Jordan, and Shamil R Sunyaev. Predicting functional effect of human missense mutations using polyphen-2. Current protocols in human genetics, 76(1):7–20, 2013.

[15] Kristian W Kaufmann, Gordon H Lemmon, Samuel L DeLuca, Jonathan H Sheehan, and Jens Meiler. Practically useful: what the rosetta protein modeling suite can do for you. Biochemistry, 49(14):2987–2998, 2010.

[16] Predrag Radivojac, Wyatt T Clark, Tal Ronnen Oron, Alexandra M Schnoes, Tobias Wittkop, Artem Sokolov, Kiley Graim, Christopher Funk, Karin Verspoor, Asa Ben-Hur, et al. A large-scale evaluation of computational protein function prediction. Nature methods, 10(3):221–227, 2013.

[17] Debora S Marks, Lucy J Colwell, Robert Sheridan, Thomas A Hopf, Andrea Pagnani, Riccardo Zecchina, and Chris Sander. Protein 3d structure computed from evolutionary sequence variation. PloS one, 6(12):e28766, 2011.

[18] Johannes Söding, Andreas Biegert, and Andrei N Lupas. The hhpred interactive server for protein homology detection and structure prediction. Nucleic acids research, 33(suppl_2):W244–W248, 2005.

[19] Andrew Waterhouse, Martino Bertoni, Stefan Bienert, Gabriel Studer, Gerardo Tauriello, Rafal Gumienny, Florian T Heer, Tjaart A P de Beer, Christine Rempfer, Lorenza Bordoli, et al. Swiss-model: homology modelling of protein structures and complexes. Nucleic acids research, 46(W1):W296–W303, 2018.

[20] Erika L McCallister, Eric Alm, and David Baker. Critical role of *β*-hairpin formation in protein g folding. Nature structural biology, 7(8):669–673, 2000.

[21] Grey W Wilburn and Sean R Eddy. Remote homology search with hidden potts models. BioRxiv, 2020.

[22] Christian A Cumbaa and Igor Jurisica. Protein crystallization analysis on the world community grid. Journal of structural and functional genomics, 11(1):61–69, 2010.

[23] Lars Malmström, Michael Riffle, Charlie EM Strauss, Dylan Chivian, Trisha N Davis, Richard Bonneau, and David Baker. Superfamily assignments for the yeast proteome through integration of structure prediction with the gene ontology. PLoS Biol, 5(4):e76, 2007.

[24] Arthur Mensch and Mathieu Blondel. Differentiable dynamic programming for structured prediction and attention. 35th International Conference on Machine Learning, ICML 2018, 8:5540–5562, 2018.

[25] Richard Durbin, Sean R Eddy, Anders Krogh, and Graeme Mitchison. Biological sequence analysis: probabilistic models of proteins and nucleic acids. Cambridge university press, 1998.

[26] Robert C Edgar. Muscle: multiple sequence alignment with high accuracy and high throughput. Nucleic acids research, 32(5):1792–1797, 2004.

[27] Cédric Notredame, Desmond G Higgins, and Jaap Heringa. T-coffee: A novel method for fast and accurate multiple sequence alignment. Journal of molecular biology, 302(1):205–217, 2000.

[28] Jianhua Zhu and Zhiping Weng. FAST: A novel protein structure alignment algorithm. Proteins: Structure, Function and Genetics, 58(3):618–627, 2005.

[29] Alexander Rives, Siddharth Goyal, Joshua Meier, Demi Guo, Myle Ott, C Lawrence Zitnick, Jerry Ma, and Rob Fergus. Biological structure and function emerge from scaling unsupervised learning to 250 million protein sequences. bioRxiv, page 622803, 2019.

[30] Michael Heinzinger, Ahmed Elnaggar, Yu Wang, Christian Dallago, Dmitrii Nechaev, Florian Matthes, and Burkhard Rost. Modeling aspects of the language of life through transfer-learning protein sequences. BMC bioinformatics, 20(1):723, 2019.

[31] Roshan Rao, Nicholas Bhattacharya, Neil Thomas, Yan Duan, Peter Chen, John Canny, Pieter Abbeel, and Yun Song. Evaluating protein transfer learning with tape. In Advances in Neural Information Processing Systems, pages 9689–9701, 2019.

[32] Ethan C Alley, Grigory Khimulya, Surojit Biswas, Mohammed AlQuraishi, and George M Church. Unified rational protein engineering with sequence-based deep representation learning. Nature methods, 16(12):1315–1322, 2019.

[33] Ahmed Elnaggar, Michael Heinzinger, Christian Dallago, Ghalia Rihawi, Yu Wang, Llion Jones, Tom Gibbs, Tamas Feher, Christoph Angerer, Debsindhu Bhowmik, et al. Prottrans: Towards cracking the language of life’s code through self-supervised deep learning and high performance computing. arXiv preprint arXiv:2007.06225, 2020.

[34] Amy X Lu, Haoran Zhang, Marzyeh Ghassemi, and Alan Moses. Self-supervised contrastive learning of protein representations by mutual information maximization. bioRxiv, 2020.

[35] Adam J Riesselman, Jung-Eun Shin, Aaron W Kollasch, Conor McMahon, Elana Simon, Chris Sander, Aashish Manglik, Andrew C Kruse, and Debora S Marks. Accelerating protein design using autoregressive generative models. bioRxiv, page 757252, 2019.

[36] Marco Cuturi and Mathieu Blondel. Soft-dtw: a differentiable loss function for time-series. arXiv preprint arXiv:1703.01541, 2017.

[37] Satoshi Koide, Keisuke Kawano, and Takuro Kutsuna. Neural edit operations for biological sequences. Advances in Neural Information Processing Systems, 2018-Decem(NeurIPS):4960–4970, 2018.

[38] Evgenii Ofitserov, Vasily Tsvetkov, and Vadim Nazarov. Soft edit distance for differentiable comparison of symbolic sequences. 2019.

[39] Svetlin A. Manavski and Giorgio Valle. CUDA compatible GPU cards as efficient hardware accelerators for Smith-Waterman sequence alignment. BMC Bioinformatics, 9(SUPPL. 2):1–9, 2008.

[40] Helen Berman, Kim Henrick, Haruki Nakamura, and John L Markley. The worldwide protein data bank (wwpdb): ensuring a single, uniform archive of pdb data. Nucleic acids research, 35(suppl_1):D301–D303, 2007.

[41] Yang Zhang and Jeffrey Skolnick. TM-align: A protein structure alignment algorithm based on the TM-score. Nucleic Acids Research, 33(7):2302–2309, 2005.

[42] Andreas Prlicć, Spencer Bliven, Peter W Rose, Wolfgang F Bluhm, Chris Bizon, Adam Godzik, and Philip E Bourne. Pre-calculated protein structure alignments at the rcsb pdb website. Bioinformatics, 26(23):2983–2985, 2010.

[43] Antonina Andreeva, Eugene Kulesha, Julian Gough, and Alexey G Murzin. The SCOP database in 2020: expanded classification of representative family and superfamily domains of known protein structures. Nucleic Acids Research, 48(D1):D376–D382, 11 2019.

[44] Christine A Orengo, Alex D Michie, Susan Jones, David T Jones, Mark B Swindells, and Janet M Thornton. Cath–a hierarchic classification of protein domain structures. Structure, 5(8):1093–1109, 1997.

[45] John Moult, Krzysztof Fidelis, Andriy Kryshtafovych, Torsten Schwede, and Anna Tramontano. Critical assessment of methods of protein structure prediction (casp)—round xii. Proteins: Structure, Function, and Bioinformatics, 86:7–15, 2018.

[46] Hua Cheng, Bong Hyun Kim, and Nick V. Grishin. MALISAM: A database of structurally analogous motifs in proteins. Nucleic Acids Research, 36(SUPPL. 1):211–217, 2008.

[47] Hua Cheng, Bong Hyun Kim, and Nick V. Grishin. MALIDUP: A database of manually constructed structure alignments for duplicated domain pairs. Proteins: Structure, Function and Genetics, 70(4):1162–1166, 2008.

[48] Cyrus Chothia, Jir Novotn, Robert Bruccoleri, and Martin Karplus. Domain association in immunoglobulin molecules. Journal of molecular biology, 186(3):651–663, 1985.

[49] Andrew W Senior, Richard Evans, John Jumper, James Kirkpatrick, Laurent Sifre, Tim Green, Chongli Qin, Augustin Žídek, Alexander WR Nelson, Alex Bridgland, et al. Improved protein structure prediction using potentials from deep learning. Nature, 577(7792):706–710, 2020.

[50] Sergey Ovchinnikov, Hetunandan Kamisetty, and David Baker. Robust and accurate prediction of residue–residue interactions across protein interfaces using evolutionary information. Elife, 3:e02030, 2014.

[51] Joe G Greener, Shaun M Kandathil, and David T Jones. Deep learning extends de novo protein modelling coverage of genomes using iteratively predicted structural constraints. Nature communications, 10(1):1–13, 2019.

[52] Adam Paszke, Sam Gross, Francisco Massa, Adam Lerer, James Bradbury, Gregory Chanan, Trevor Killeen, Zeming Lin, Natalia Gimelshein, Luca Antiga, et al. Pytorch: An imperative style, high-performance deep learning library. In Advances in neural information processing systems, pages 8026–8037, 2019.

[53] WA Falcon. Pytorch lightning. GitHub. Note: https://github.com/PyTorchLightning/pytorch-lightning, 3, 2019.

[54] Peter JA Cock, Tiago Antao, Jeffrey T Chang, Brad A Chapman, Cymon J Cox, Andrew Dalke, Iddo Friedberg, Thomas Hamelryck, Frank Kauff, Bartek Wilczynski, et al. Biopython: freely available python tools for computational molecular biology and bioinformatics. Bioinformatics, 25(11):1422–1423, 2009.

[55] John D Hunter. Matplotlib: A 2d graphics environment. Computing in science & engineering, 9(3):90–95, 2007.

[56] Pauli Virtanen, Ralf Gommers, Travis E Oliphant, Matt Haberland, Tyler Reddy, David Cournapeau, Evgeni Burovski, Pearu Peterson, Warren Weckesser, Jonathan Bright, et al. Scipy 1.0: fundamental algorithms for scientific computing in python. Nature methods, 17(3):261–272, 2020.

[57] Charles R Harris, K Jarrod Millman, Stéfan J van der Walt, Ralf Gommers, Pauli Virtanen, David Cournapeau, Eric Wieser, Julian Taylor, Sebastian Berg, Nathaniel J Smith, et al. Array programming with numpy. arXiv preprint arXiv:2006.10256, 2020.

